# Hessian single molecule localization microscopy using sCMOS camera

**DOI:** 10.1101/385112

**Authors:** Fudong Xue, Wenting He, Fan Xu, Mingshu Zhang, Liangyi Chen, Pingyong Xu

## Abstract

Single-molecule localization microscopy (SMLM) has the highest spatial resolution among the existing super-resolution (SR) imaging techniques, but its temporal resolution needs further improvement. An sCMOS camera can effectively increase the imaging rate due to its large field of view and fast imaging speed. Using an sCMOS camera for SMLM imaging can significantly improve the imaging time resolution, but the unique single pixel-dependent readout noise of sCMOS cameras severely limits their application in SMLM imaging. This paper develops a Hessian-based SMLM (Hessian-SMLM) method that can correct the variance, gain and offset of a single pixel of a camera and effectively eliminate the pixel-dependent readout noise of sCMOS cameras, especially when the signal-to-noise ratio is low. Using Hessian SMLM to image mEos3.2-labeled actin was able to significantly reduce the artifacts due to camera noise.

## Introduction

Super-resolution (SR) microscopy enables biological researchers to see nanoscale images of intracellular structures. Single-molecule localization microscopy (SMLM), such as PALM/STORM(Betzig et al, 2006; Hell, 2007; Rust et al, 2006), use photocontrollable fluorescent proteins to label target proteins and image a small number of single molecules that are randomly activated at different times. And then the central position of a single fluorescent molecule is precisely located by a Gaussian fitting algorithm. Next, a final image with diffraction-unlimited resolution is reconstructed if a sufficient number of single molecules are obtained. This SMLM technology can achieve a spatial resolution of approximately 20-30 nanometers. However, this method requires thousands of image frames to reconstruct SR structures, and it is difficult to meet the researchers’ requirements for temporal resolution.

To achieve higher temporal resolution, a faster signal acquisition with a sufficiently high signal-to-noise ratio is important. Increasing the exposure frequency (i.e., decreasing the exposure time) provides faster signal acquisition, but for back-illuminated electron multiplying charge-coupled devices (EMCCDs), the read speed is currently limited to 70 frames per second, limiting the temporal resolution of SR techniques that use multiple sequential frames to reconstruct a diffraction-unlimited image, especially SMLM microscopy. Compared with EMCCD cameras, the newly developed sCMOS camera with the ability to detect single molecules has the advantages of a large field of view, fast readout speed, etc., which can significantly increase the data acquisition rate and improve the temporal resolution. However, unlike an EMCCD, an sCMOS camera has a unique single-pixel-dependent readout noise, and the random noise appearing in each single frame at different time points will be incorrectly fitted to the fluorescence signal of a single molecule. Since an SMLM image is a superposition of a single molecule localized in a large number of single-frame (tens of thousands to several tens of thousands) images, the superposition of randomly occurring noise in a single-frame image generates a pseudostructure in the final SR image.

The unique noise of the sCMOS camera limits its application in fast SMLM imaging. Therefore, there is an urgent need for noise calibration and for the development of a corresponding single-molecule localization algorithm. Huang et al developed an algorithm to calibrate the independent readout noise by addressing three important characteristics (Huang et al, 2013): 1) Remove single-pixel-dependent offsets; 2) Correct single-pixel-dependent gain values; and 3) Remove the systematic fluctuation noise of each pixel. This method first calculates the brightness offset and the fluctuation level of each pixel by acquiring the dark image of the camera and then calculates the gain per pixel by illuminating the camera with different intensities. Then, these three parameters are used to calibrate the actual fluorescence image. This method is a good way to distinguish the source of readout noise and to calibrate and eliminate the single-pixel-dependent systematic readout noise of the camera. However, the Huang et al, assumed Gaussian noise model of readout noise for all pixels on the sensor, which is based on the data obtained when the camera is dark, may not reflect the underlying fluctuation pattern for each pixel on the camera at different time points during the actual sampling, because during the actual sampling the fluctuation level of each pixel at a certain time point varies greatly. In SMLM experiments, these fluctuating noises can easily be treated as signals by single-molecule detection algorithms, resulting in artifacts in the reconstruction results.

The Hessian matrix is a square matrix composed of second-order partial derivatives of multivariate functions and is often used in the field of boundary detection and denoising (Lefkimmiatis et al, 2013; Sun et al, 2015). In 2018, Huang X. et al developed a Hessian-SIM method (Huang et al, 2018). The Hessian penalty matrix was used to remove the scattering noise after SIM reconstruction, and the SR images with minimal artifacts were obtained. In this paper, we develop the new technique Hessian SMLM, an SMLM imaging technology for rapid imaging using an sCMOS camera and combining the Hessian and Huang’s sCMOS calibrating algorithms to effectively remove sCMOS pixel-dependent readout noise. The new method can significantly reduce the reconstruction artifacts due to noise when used for actin PALM imaging.

## Overview of Hessian-SMLM for SR imaging

First, we collected the image when the sCMOS camera was in the dark or illuminated by a series of white lights with different intensities and obtained the calibration parameters of the sCMOS camera: Offset and Gain. Then, a fluorescently labeled biological sample was imaged by the sCMOS camera. The corresponding pixel offset obtained previously was subtracted from each pixel in the acquired single-molecule data, and the fluorescence of each pixel was then divided by the Gain. A Hessian denoising algorithm was then used to remove the fluctuation noise caused by Variance. Next, the noise-corrected single-molecule image was used for single-molecule extraction. Last, the precise single-molecule position was obtained by Gaussian fitting, and the SR image was reconstructed. The specific process of Hessian-SMLM is shown in Figure 1.

**Figure 1.**
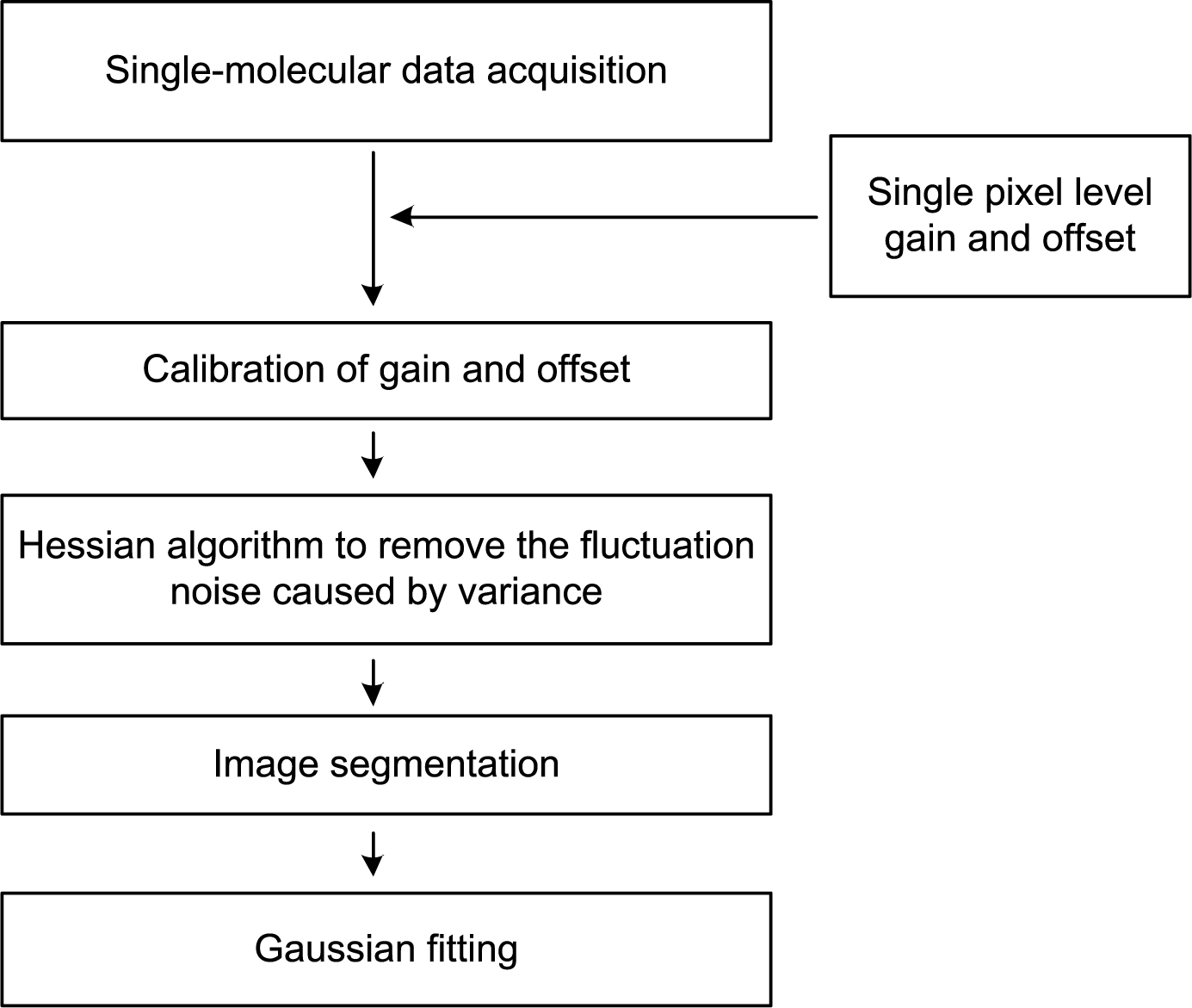
Diagram of Hessian SMLM SR imaging. First, single pixel level characterization of sCMOS is obtained, and the single-molecule data of each pixel are calibrated by the gain and offset values of the sCMOS camera. Then, the calibrated data are treated with the Hessian algorithm to remove the noise caused by variance. In the localization step, the images are segmented and fitted by Gaussian fitting to obtain the final SR image.

### 1. Acquisition of offset and gain of sCMOS camera

Offset is a constant luminance term in the readout process of sCMOS that is added in advance to avoid the negative luminance caused by the readout noise in the sCMOS camera. By taking a series of dark images with an sCMOS camera and calculating the average value of each pixel, the offset of each pixel can be measured. First, we turned off the autocalibration mode of the Prime 95B sCMOS camera (Photometrics, Tucson, AZ USA) and recorded 60,000 frames of dark images (Figure 2a). The offset of each pixel in the sCMOS camera can be derived from the following statistical analysis

**Figure 2.**
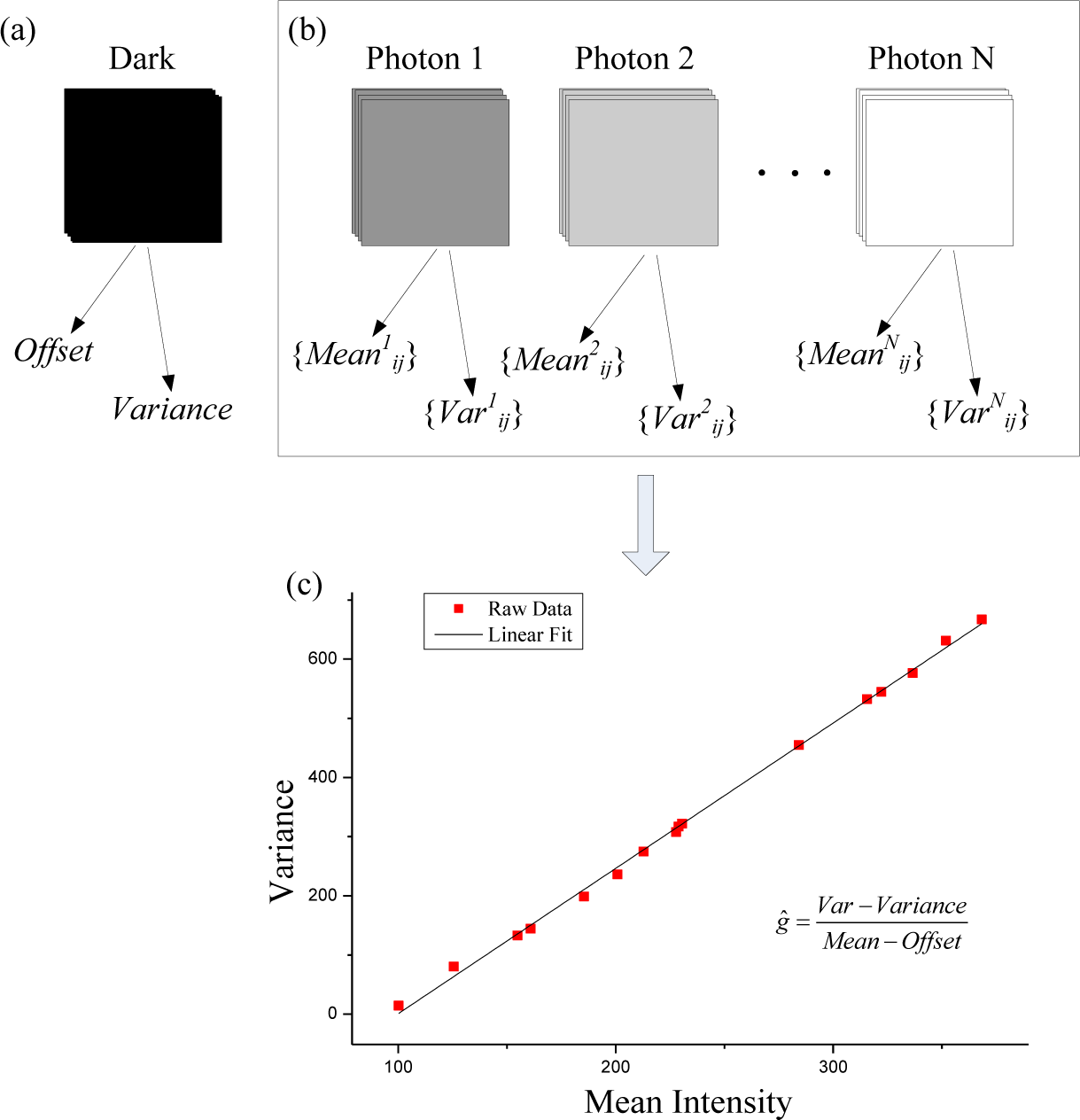
sCMOS characterization at the single pixel level. (a) To measure the offset and variance of sCMOS, 60,000 dark frames were used. (b) A series of image sequences were recorded at different average intensity levels to measure the mean values and variances. (c) The gain for each pixel can then be calculated with the help of the previously obtained variance and mean value. The red dashed line is the variance at different mean intensities of one pixel of the sCMOS camera, and the solid line shows the linear fit result.

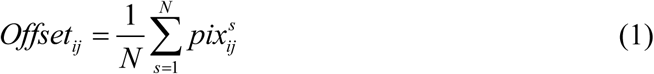

Where *N* is the total number of dark frames and 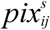 is the photon number at frame *s* for pixel *ij*.

Because the gain of each pixel of the sCMOS camera is different, the amplification of the fluorescence signal is different for each pixel during actual imaging, so it is necessary to measure the gain of each pixel for calibration. To calculate the gain for each pixel, we first took a series of dark images and calculate the variance as follows

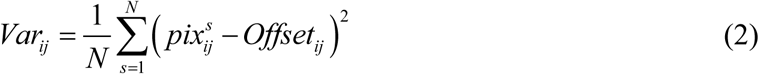

Then, we uniformly illuminated the camera with white light of different light intensities ranging from ~20 to 200 photons, which were evenly divided into 15 groups from low to high according to the light intensity. In total, 20,000 frames of images were captured for each group (Figure 2b). The variance and mean value of brightness under different light intensities were then obtained, and linear fitting by the least-squares method was performed, as in equation (3) (Figure 2c).

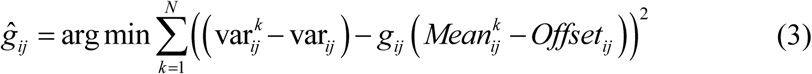

### 2. Hessian algorithm removes variance noise in single-molecule images

For Prime 95B sCMOS imaging, the pixel size is 110 nm/pixel for a 100× magnification microscope. Due to the optical diffraction limit, the full width at half maximum (FWHM) of a diffraction spot is greater than 250 nm, so when a single molecule on the sample is imaged through an optical system, the molecule occupies at least two pixels on the camera, which means that spatial correlation exists for two pixels. At the same time, if the speed of single-molecule blinking is lower than the camera’s readout speed, there should also be correlations across time series for the same pixel. However, there should be no correlation of random Gaussian noise, especially sCMOS pixel-dependent noise, between different pixels and/or across time series for the same pixel. Therefore, we refer to a reference (Huang et al, 2018) and introduce the Hessian penalty to remove the temporally and spatially random noise. The optimization function is as follows

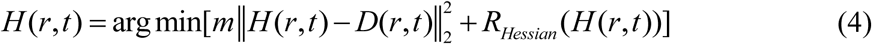

where *D* is the raw data collected by the camera, *H* is the optimized data, *m* is the relative weight between the first term and the Hessian penalty, and *R_Hessian_* is the Hessian penalty(Lefkimmiatis et al, 2012), which is expressed by formula (5)

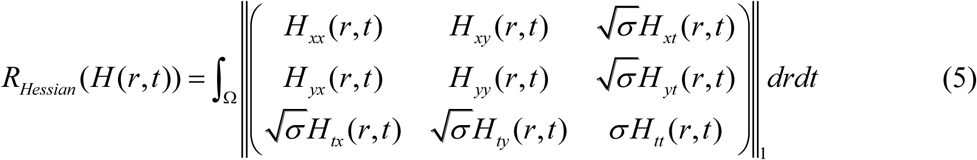

where *r* is the position of each pixel, Ω represents the integral area that contains all pixels within the image *H*, and *H_xy_* is the second-order partial derivative of *H* versus *xy*, σ is a parameter that was introduced to enforce the continuity of structures along the time axis.

We thus finally obtain the optimal noise-removed single-molecule data by calculating the minimum value of formula (4) from the single-molecule data, and use these data for single molecule extraction and reconstruction as previous reference (Olivo-Marin, 2002; Smith et al, 2010).

### 3. Cell culture

U2OS cells were cultured using complete MCMM (McCoy’s 5A Medium Modified, Gibco) supplemented with 10% inactivated fetal bovine serum (FBS, Gibco) in a 37°C incubator with 5% CO_2_. The construct of LifeAct-mEos3.2 for labeling actin structures and the Hessian-SMLM SR imaging were described in (Zhang et al, 2012). The constructs were transfected using Lipofectamine™ 2000 Transfection Reagent (Invitrogen, USA). After a 48-h transfection, the cells were then fixed by 4% (w/v) paraformaldehyde (NOVON Scientific, Pleasanton, CA USA) and 0.2% glutaraldehyde (Electron Microscopy Sciences, Hatfield, PA USA) and imaged by a custom built PALM microscope.

### 4. Simulation of Hessian SMLM imaging

To verify the Hessian SMLM’s ability to remove variance noise, we simulated single-molecule data containing fluctuating noise and performed PALM imaging simulations. We simulated and generated the original data shown in Figure 3b and used the Thunder STORM imaging plug-in to generate single-molecule images (Ovesný et al, 2014). Simulated wave noise was added (Figure 3a) so that we obtained simulated single-molecule data containing fluctuating noise. Then, the PALM single-molecule localization algorithm was used to reconstruct the image with (Figure 3d) and without (Figure 3c) Hessian denoising methods. As shown in Figure 3c,d, the resulting reconstruction without the Hessian denoising method contains more artifacts, while Hessian denoising efficiently removed the artifacts caused by the fluctuating noise.

**Figure 3.**
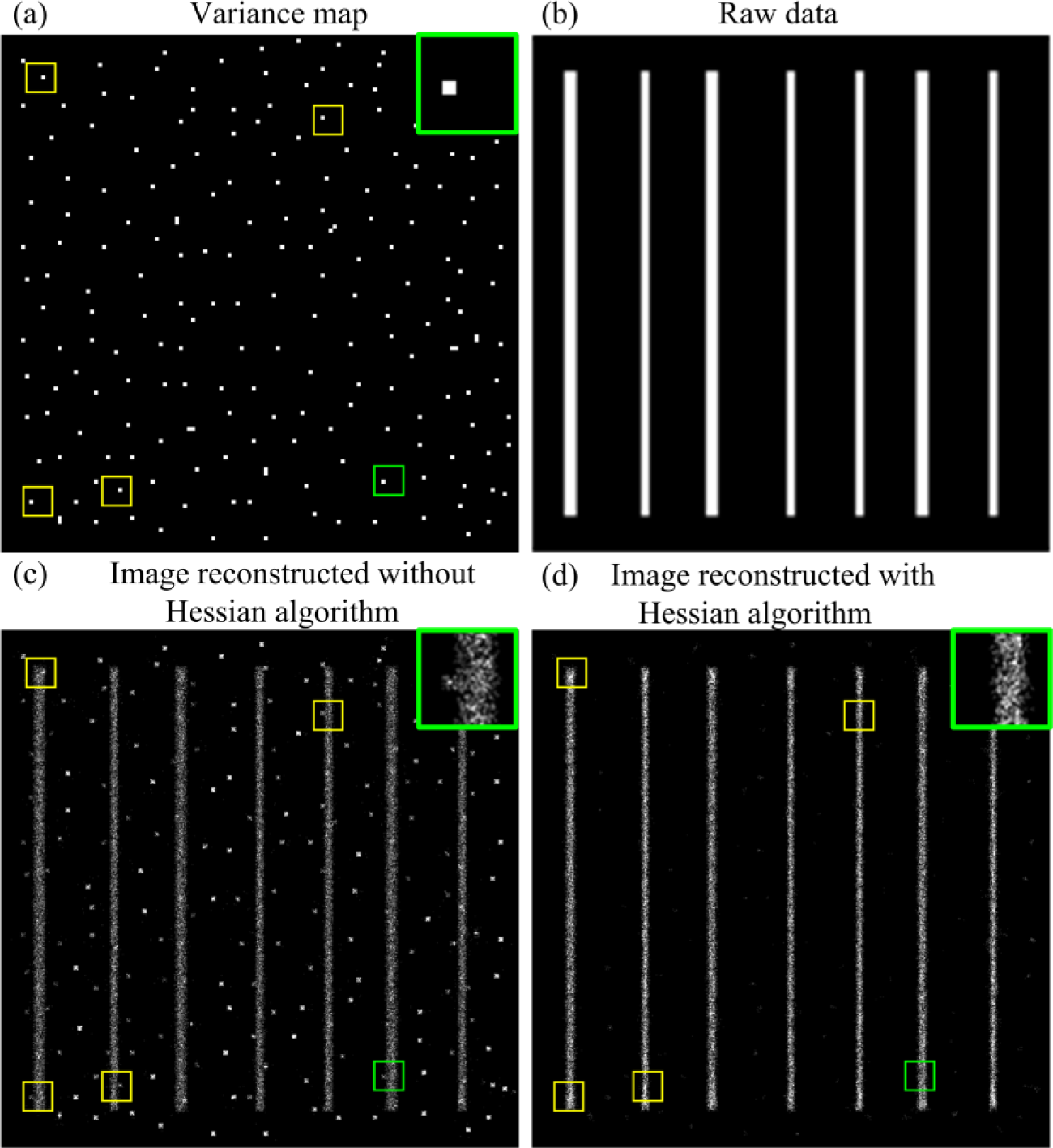
Simulation of single emitters on parallel lines based on the shown variance map with localization using single-molecule localization microscopy with and without the Hessian algorithm. (a) Simulated variance map. (b) Raw data used to simulate the single-molecule image. (c, d) Simulated single-molecule image reconstructed without (c) and with (d) the Hessian algorithm. The green box denotes the enlarged section shown in the inset.

### 5. Hessian SMLM imaging of actin

To verify the effectiveness of Hessian SMLM in SR imaging of biological samples, we transfected LifeAct-mEos3.2 in U2OS cells and acquired single-molecule data using the Prime 95B sCMOS camera. We treated the single-molecule data to reconstruct an SR image using different algorithms separately as follows: 1). Single-molecule localization algorithm of PALM; 2). The sCMOS-calibrated single-molecule localization algorithm by Huang et al; and 3). The Hessian SMLM algorithm in the current study.

We found that when using the Prime 95B CMOS camera to acquire single-molecule data, there is substantial noise in the variance map of the fluorescence signal. Most of the noise is completely colocalized with the variance map of the dark camera (Figure 4c), suggesting that the noise comes mainly from the variance when the camera is dark. Moreover, when the single-molecule data collected by the sCMOS camera are directly used in the PALM single-molecule localization algorithm, the reconstruction results contain more artifacts (Figure 4d, indicated by boxes).

**Figure 4.**
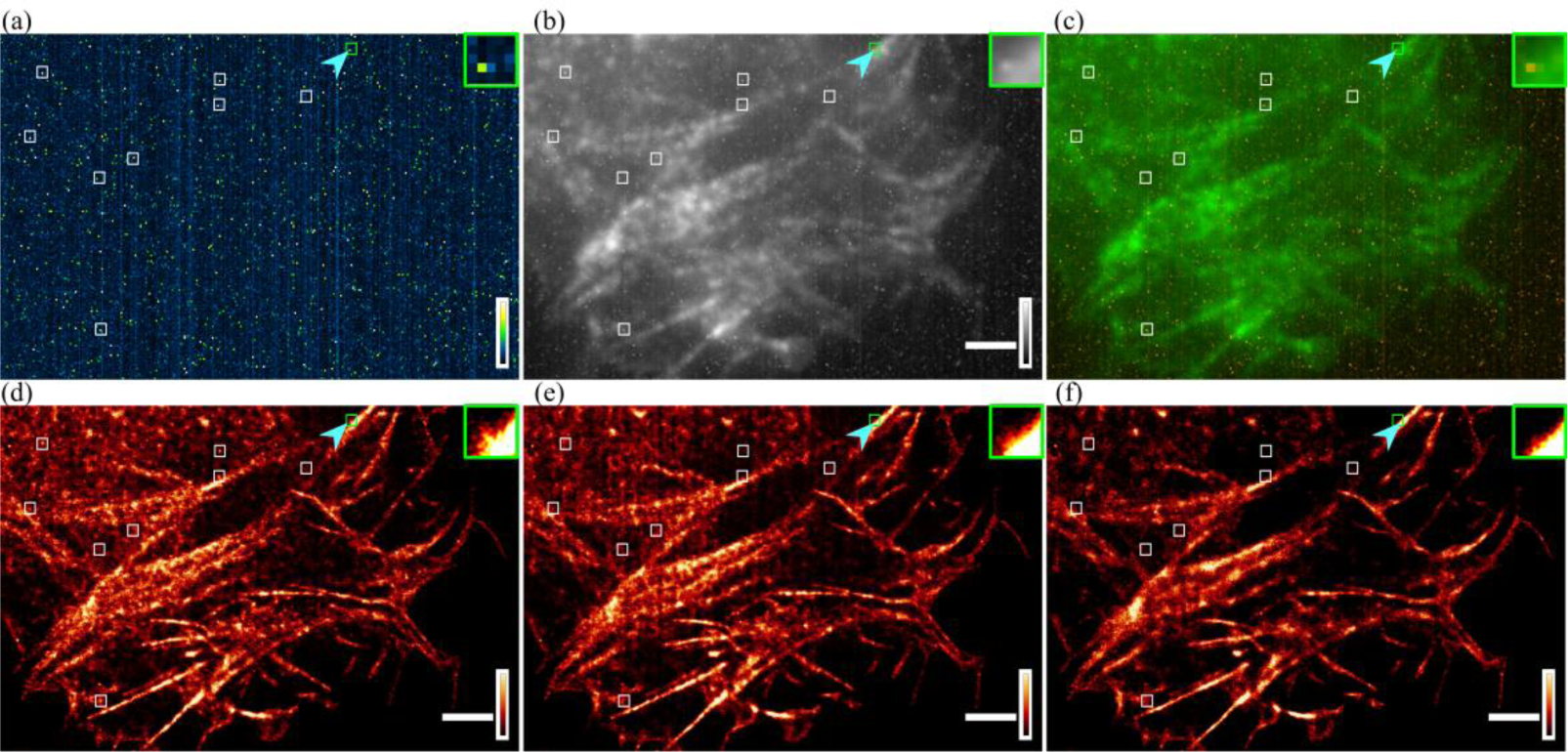
Hessian SMLM imaging of actin with three SMLM algorithms. (a) Readout variance map of the sCMOS camera. (b) Standard deviation (STD) projection image of single-molecule data. (c) Merged image of the variance map (a) and the STD projection image (b). Green: pixel-level readout variance of the dark camera; Red: STD image. (d-f) Reconstructed super-resolution images of actin analyzed using the conventional algorithm (d), sCMOS camera-specific algorithm (e) and Hessian SMLM algorithm (f). The green box denotes the enlarged section shown in the inset. Color scales, bottom to top: 0-65 ADU^2^, where ADU = analog-to-digital units (a); minimum-to-maximum signal (b); same upper bound chosen for best visualization for d-f. Scale bar: 2 μm.

Obviously, these artifacts come from the variance of the sCMOS camera because they are exactly the same noise points in the variance map when the camera is dark (Figure 4a). Using Huang et al’s sCMOS correction method to correct the variance, gain, and offset greatly improved the reconstruction results and significantly reduced the artifacts due to single-pixel fluctuations. However, for some areas, especially those where the variance is smaller, the calibration effect of this method is not ideal (Figure 4e). One possible reason is that the variance we obtained with the dark camera can only represent the statistical fluctuation level of each pixel during the sampling process. In the actual sampling, the averaged variance of each pixel from the dark sCMOS camera does not accurately characterize the true fluctuation level of each frame at different time points. However, as shown in Figure 4f, Hessian SMLM can efficiently remove the variance noise in each frame produced by the fluctuation at different time points, and the artifacts generated in the reconstruction result are significantly reduced.

Next, we further compared the effect of the above three methods on a LifeAct-mEos3.2 reconstruction with a low exposure time and laser intensity. The low exposure time and laser intensity significantly reduced the signal-to-noise ratio of the single-molecule data. The structure directly reconstructed with the PALM localization algorithm had substantial noise and many artifactual structures (Figure 5b). Using the sCMOS calibration method removed some of the noise (Figure 5c), but the effect was not as good as the effect when the signal-to-noise ratio was high (Figure 4e). However, with the Hessian SMLM method, the reconstruction artifacts were basically removed, and the structural continuity of actin was much better than the continuity with the other two methods (Figure 5d compared to 5b and 5c, and enlarged regions shown in Figure 5e). We chose four pixels with different variance and measured the pixel intensity before and after Hessian algorithm (Figure 5f). The fluctuating value of the four pixels decreased significantly after Hessian algorithm (Figure5f, blue and red solid lines), while the mean pixel values are almost the same before and after Hessian algorithm (Figure5f, blue and red dash lines). Therefore, Hessian SMLM is able to efficiently remove the variance noise of each frame produced by the fluctuation at different time points, and reduce the artifacts.

**Figure 5.**
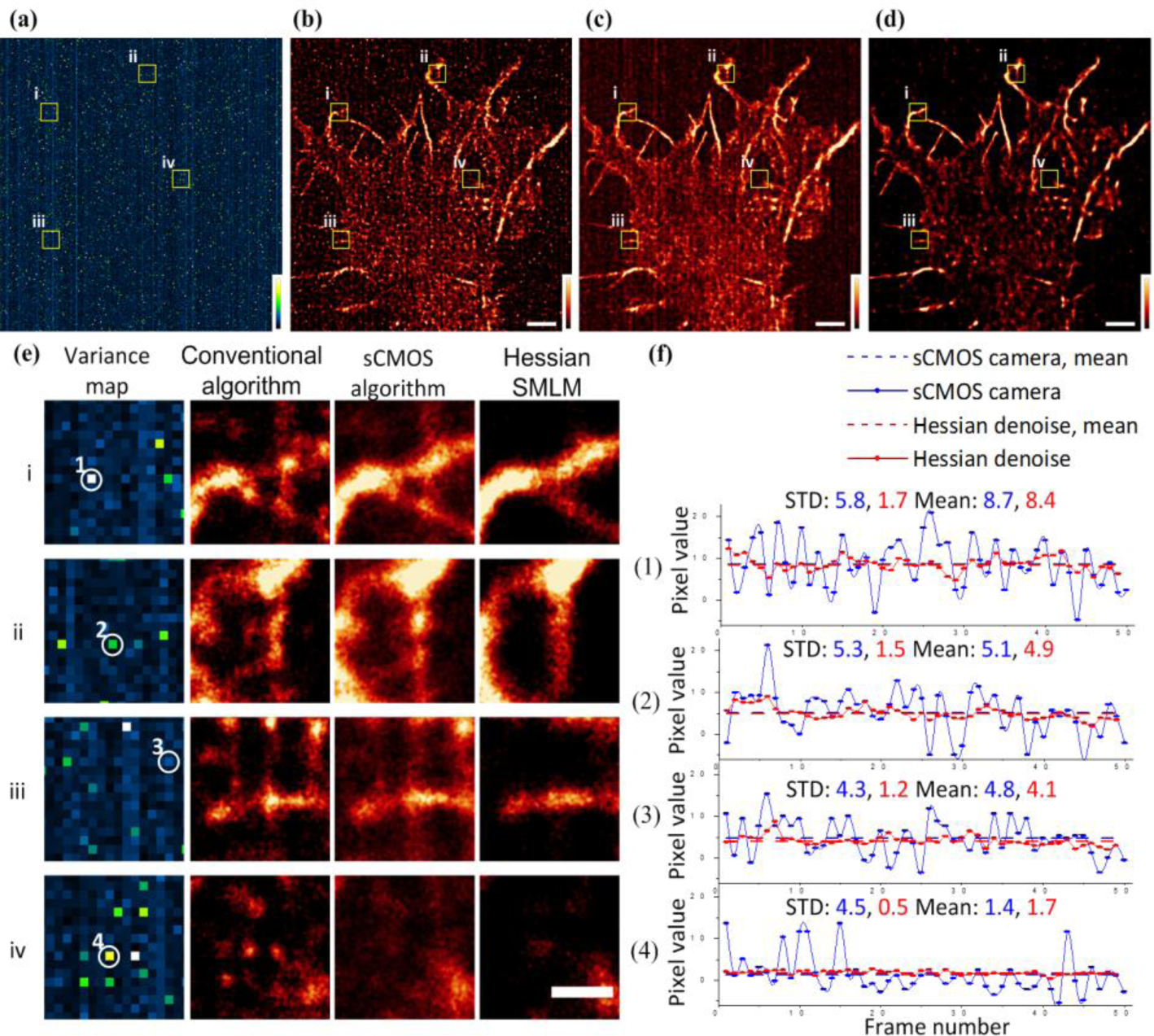
Hessian SMLM significantly removes the noise of sCMOS and artifacts in SMLM images even when the signal-to-noise ratio is low. (a) Readout variance map of the sCMOS camera. (b-d) Reconstructed super-resolution images of actins analyzed using the conventional algorithm (b), sCMOS algorithm (c) and Hessian SMLM algorithm (d). (e) Zoom in regions (i-iv) from (a-d) show actin structures by the PALM localization algorithm, the sCMOS calibration method and Hessian SMLM algorithm. (f) The pixel intensity traces of the selected pixels were from cropped regions (i-iv) over 50 frames. Color scales, bottom to top: 0-50 ADU^2^ (a); same upper bound chosen for best visualization for b-d. Scale bars: (a-d) 2 μm, (e) 0.2 μm.

## Discussion

Although sCMOS cameras are inferior to EMCCDs for single-molecule imaging, sCMOS cameras have the advantages of fast speed and large field of view and have potential applications in live cell SMLM imaging. However, the unique single-pixel-dependent readout noise of sCMOS cameras severely limits their application in live SMLM, especially when the signal-to-noise ratio of a single-molecule signal is low. For SMLM, there are at least two situations that produce low signal-to-noise ratios for single-molecule signals. The first situation is the imaging of fixed cells by SMLM when the signal-to-noise ratio becomes worse as the sampling time increases. The other one is live SMLM. For live SMLM, on the one hand, the exposure time must be decreased to increase the sampling frequency and thus improve the temporal resolution. On the other hand, it is necessary to reduce the irradiation light intensity to reduce phototoxicity. However, reducing the exposure time and reducing the intensity of the illumination light significantly reduce the signal-to-noise ratio of the fluorescence signal. Under these conditions, the camera’s readout noise will be particularly notable. In particular, the camera’s variance noise will be Gaussian fitted as a fake single-molecule signal, which will produce false structures or artifacts in the final reconstructed SR image. In this study, our new Hessian SMLM SR imaging method can well remove the single-pixel-dependent readout noise and effectively reduce the artifacts caused by the readout noise of the sCMOS camera. This method is especially suitable for SMLM imaging when the signal-to-noise ratio is low and has greater advantages over other SMLM imaging methods under conditions of low exposure time and laser intensity. Therefore, Hessian SMLM is potentially more suitable than the other methods for live cell SMLM imaging.

## ACKNOWLEDGEMENT

This project was supported by the National Key R&D Program of China (2016YFA0501500 and 2017YFA0505300), the National Natural Science Foundation of China (31421002, 21778069 and 31670870), Project of the Chinese Academy of Sciences (XDB08030203 and CAS-Peking University Joint Team Project). We thank Dr. Fang Huang from Purdue University for very helpful comments on the manuscript.

